# Characterization of Human Endogenous Retroviruses in Gastric Cancer with *Helicobacter pylori*: A Study from Northern Brazil

**DOI:** 10.1101/2025.09.06.674554

**Authors:** Marcos da Conceição, Juliana Barreto Albuquerque Pinto, Diego Pereira, Rubem Ferreira da Silva, Sérgio Augusto Antunes Ramos, Daniel de Souza Avelar, Jéssica Manoelli Costa da Silva, Ronald Matheus da Silva Mourão, Valéria Cristiane Santos da Silva, Samir Mansour Casseb, Samia Demachki, Tatiane Neotti, Ana Karyssa Mendes Anaissi, Williams Fernandes Barra, Amanda Ferreira Vidal, Geraldo Ishak, Paulo Pimentel de Assumpção, Rommel Mario Rodriguez Burbano, Fabiano Cordeiro Moreira

## Abstract

Human endogenous retroviruses (HERVs) are retroelements that have integrated their genetic material into the human genome, accumulating mutations over time and accounting for approximately 8% of the genome. Under abnormal deregulation conditions, these elements can be expressed and contribute to the development of diseases, such as gastric cancer. This malignancy may be associated with infections, including those caused by *Helicobacter pylori*. However, the scientific literature does not yet provide clear evidence regarding the relationship between HERVs and *H. pylori* in the context of gastric cancer. Thus, HERVs may represent potential biomarkers for this neoplasm, as well as possible therapeutic targets. This study aimed to characterize HERV expression in gastric cancer using next-generation sequencing (NGS). We analyzed 46 tumor tissue samples and 42 peritumoral tissue samples from patients diagnosed with gastric adenocarcinoma, collected at HUJBB and Ophir Loyola hospitals. Among the tumor samples, 38 tested positive for H. pylori infection. For library preparation, 1 μg of total RNA per sample was used, with integrity assessed via TapeStation (∼260 bp band). cDNA libraries were sequenced using the Illumina NextSeq 500 platform (paired-end), following the ID Output V2 kit protocol. Alignment was performed with STAR software, and HERVs were identified and quantified using Telescope. Differential expression analysis of HERVs was performed on transcript data using DESeq2. A total of 183 HERVs were found to be differentially expressed in tumor tissues compared to adjacent tissues. In tumor samples associated with *H. pylori* infection, 44 HERVs showed differential expression. Overall, tumor tissues exhibited higher HERV transcription compared to adjacent tissues.

## INTRODUCTION

Human Endogenous Retroviruses (HERVs) are retroelements in the human genome that resemble the genetic material of exogenous retroviruses that infected the germ cells of the host species during its evolutionary history (Bao *et al*., 2024). Since then, they have become integrated into our genomes and lost the ability to produce horizontally transmitted pathogenic virions (Johnson, 2019).

It is known that HERVs contain all the necessary regulatory elements for transcription, are active only in vertebrates and constitute approximately 8% of the human genome (Geis and Goff, 2020). They present insertions with numerous accumulated mutations and deletions, although some elements have retained intact open reading frames (ORFs) capable of encoding functional protein (Srinivasachar Badarinarayan and Sauter, 2022). All HERVs represent seemingly extinct retroviral lineages, and in many normal human tissues, their gene expression can be detected. However, most of them have mutations that prevent the expression of proteins and viral particles (Burn *et al*., 2022). Some HERVs, mainly HERV-K (HML-2), have the potential for protein coding (Vargiu *et al*., 2016). When it does occur, the level and pattern of expression can be modulated under pathological conditions, such as cancer (Löwer, 1999).

In recent decades, research on infections associated with carcinogenesis has intensified, including the identification of altered cellular mechanisms aimed at developing interventions that can reduce the risk and progression of various types of tumors (Cegolon *et al*., 2013). However, studies involving human endogenous retroviruses (HERVs) with the potential to induce malignancies remain limited (Kitsou, Lagiou, and Magiorkinis, 2023).

Stomach cancer is among the most significant malignancies, with over 960,000 new cases reported in 2022, ranking as the fifth most common and the fifth deadliest cancer worldwide (Bray *et al*., 2024). The highest incidence rates of gastric cancer are found in countries in East Asia, Central and Eastern Europe, and South and Central America (Balakrishnan *et al*., 2017). Notably, the Northern region of Brazil, particularly the city of Belém, has shown an increase in gastric cancer mortality over recent decades (Curado *et al*., 2019).

One of the pathogens potentially involved in the etiology of gastric cancer is the bacterium *Helicobacter pylori*, which colonizes the gastrointestinal mucosa of approximately half of the global population (Duan *et al*., 2025). This bacterium is also associated with other gastrointestinal diseases, including chronic gastritis, gastric ulcers, and duodenal ulcers (Li, He and Lu, 2024).

*H. pylori* is currently considered a resident bacterium of the gastric environment with significant clinical relevance. Chronic infection by this bacterium is the main risk factor for the development of non-cardia gastric cancer (IARC, 1994; Plummer *et al*., 2015). It is estimated that approximately 4.4 billion people worldwide came into contact with the bacteria, which correlates with the global incidence of gastric cancer (Hooi *et al*., 2017).

RNA sequencing (RNA-Seq), performed through next-generation sequencing (NGS) systems, is widely recognized as a powerful and cost-effective tool for transcriptome analysis, providing researchers with comprehensive insights into gene expression patterns that offer high sensitivity and resolution (Hu *et al*., 2024). Simultaneously, numerous in silico techniques have been developed for identifying and annotating potential HERVs (Lerat, 2010).

HERVs can play a significant role in cancer development, although their exact functions and mechanisms remain largely unknown. For this reason, studying these retroelements is considered promising, especially when they are linked to malignancies such as gastric cancer. Furthermore, they hold potential as important biomarkers and as targets for the development of novel therapies. Therefore, this study aims to characterize differentially expressed HERVs in gastric adenocarcinoma.

## METHODOLOGY

### Sample Characterization and Ethical Aspects

As part of the overarching project to which this proposal belongs, 46 tumor tissue samples and 42 samples of peritumoral adjacent to the tumor were collected from patients diagnosed with gastric adenocarcinoma. Of this total, 38 tumor samples tested positive for *H. pylori* infection, while 6 tested negative. The samples were obtained from the João de Barros Barreto University Hospital (HUJBB) and the Ophir Loyola Hospital.

All participants were fully informed about the research objectives, and samples were collected only after obtaining informed consent through the signing of the Free and Informed Consent Form (TCLE). The use of all samples and the execution of this study were approved by the Research Ethics Committee of the João de Barros Barreto University Hospital, under protocol number CAAE 47580121.9.0000.5634. For tumor tissue collection, 0.5 cm fragments were surgically resected. These fragments were collected immediately following gastric resection, preserved in RNAlater for transport, and stored in a -80 °C freezer.

### Clinical Characterization of the Patients

The medical records and other clinical data of the investigated patients were reviewed for the presence of gastric adenocarcinoma, as well as their gender, age, Lauren histological subtype, TNM pathological staging, administration of perioperative chemotherapy regimens with 5-Fluorouracil, Leucovorin, Oxaliplatin, and Taxane (FLOT), and mortality. TNM staging was determined based on the 8th edition of the AJCC TNM classification for gastric cancer.

### Total RNA Extraction

Initially, approximately 50–100 ng of tissue from each sample was macerated. Then, 1 mL of TRIZOL® reagent was added to the processed tissue for extraction. TRIZOL® reagent (Thermo Fisher Scientific) was used to preserve cellular RNA integrity and promote cell lysis. After centrifugation at 13,000 rpm for 10 minutes at 4 °C, RNA was recovered by precipitation with isopropyl alcohol. The precipitated RNA (total RNA) was washed with ethanol (EtOH), air-dried at room temperature, and subsequently analyzed for integrity and concentration using the Qubit 2.0 Fluorometer (Thermo Fisher Scientific), NanoDrop ND-1000 (Thermo Fisher Scientific), and 2200 TapeStation System (Agilent). Ideal criteria for total RNA integrity included values between 1.8 and 2.2 for the A260/A280 ratio, greater than 1.8 for the A260/A230 ratio, and an RNA Integrity Number (RIN) of 5 or higher. Finally, the extracted total RNA was stored in a -80 °C freezer until use.

### cDNA Library Construction

For library construction, the TruSeq Stranded Total RNA Library Prep Kit with Ribo-Zero Gold (Illumina, US) was used according to the manufacturer’s instructions. In the library preparation, 1 μg of total RNA was used per sample, in a final volume of 10 μL. Once the libraries were constructed, a new integrity assessment was performed using the 2200 TapeStation System. At the end of the process, a band of ∼260 base pairs was observed.

### NGS Sequencing and Read Quality Control

The previously constructed cDNA libraries were loaded onto the Illumina NextSeq sequencing system and sequenced using the paired-end method (reads were generated from both the forward and reverse strands of the cDNA). The NextSeq 500 ID Output V2 kit (Illumina, 150 cycles) was used to process the libraries under the conditions specified by the manufacturer. Base-called reads were converted into FASTQ format using the Reporter software, where the sequences and their quality scores were encoded in ASCII format. Subsequently, read quality was assessed using FastQC v0.12.0. Adapter sequences and low-quality reads were removed using Trimmomatic v0.39 (Bolger, Lohse and Usadel, 2014).

### Alignment

Filtered and trimmed reads from the previous step were aligned against human transcriptome sequences, using the hg38 human genome transcripts as the reference index. Alignment was performed using STAR software v2.7.10b (Dobin *et al*., 2013). Subsequently, the identification and quantification of HERV transposable elements were carried out using Telescope software v1.0.3 (Bendall *et al*., 2019), based on previously described annotations available at https://github.com/mlbendall/telescope_annotation_db/tree/master/builds. The resulting data were imported using the awk scripting language for downstream analysis in R software v4.5.1 (R Core Team, 2025). Samples with transcript counts below 1,000 were excluded from the analysis.

### Differential Analysis

To detect the abundance of differentially expressed HERVs, the DESeq2 package (version 3.14) (Love, Huber and Anders, 2014) was used. The results obtained from DESeq2 were normalized and employed in subsequent analyses. To define differentially expressed transcripts across conditions, such as tumor and adjacent tissues for all gastric cancer samples, and tumor tissues alone, a threshold of log2FoldChange > 1 and adjusted p-value < 0.05 was applied. HERV expression was visualized using the ggplot2 package (Wickham, 2016) and ComplexHeatmap v2.20.0 (Gu, 2022). The Receiver Operating Characteristic (ROC) curve was plotted using the pROC package v1.18 (Robin *et al*., 2011) to evaluate the sensitivity and specificity of each transcript in relation to the clinical feature in which it was differentially expressed, based on an Area Under the Curve (AUC) > 0.7 when comparing tumor and peritumoral tissues, and 0.8 when comparing tumor tissues with *H. pylori*-positive versus *H. pylori*-negative samples. For HERVs with favorable AUC values, Shapiro-Wilk tests were performed comparing adenocarcinoma and adjacent tissues, followed by Wilcoxon tests with p-values adjusted using the Bonferroni method.

## RESULTS

To perform the differential expression analysis of gastric cancer tumor tissues, all available adenocarcinoma and peritumoral samples were included, regardless of infection status. This analysis identified 183 transcripts with significant differential expression between tumor and adjacent tissues, of which 164 were upregulated and 19 downregulated. Additionally, a separate differential expression analysis was conducted on adenocarcinoma tissues, comparing *H. pylori*-positive and *H. pylori*-negative samples, which revealed 44 transcripts with significant changes - 38 downregulated and 6 upregulated.

Next, the AUC metric was applied to filter HERVs with the highest discriminatory power between case and control samples. As a result, 12 HERVs were selected, belonging to the families ERVLE, HERVL, HERVL66, HERVW, HML2, HML3, MER101 and MER61 (Fig. 1a).

**Fig. 1.**
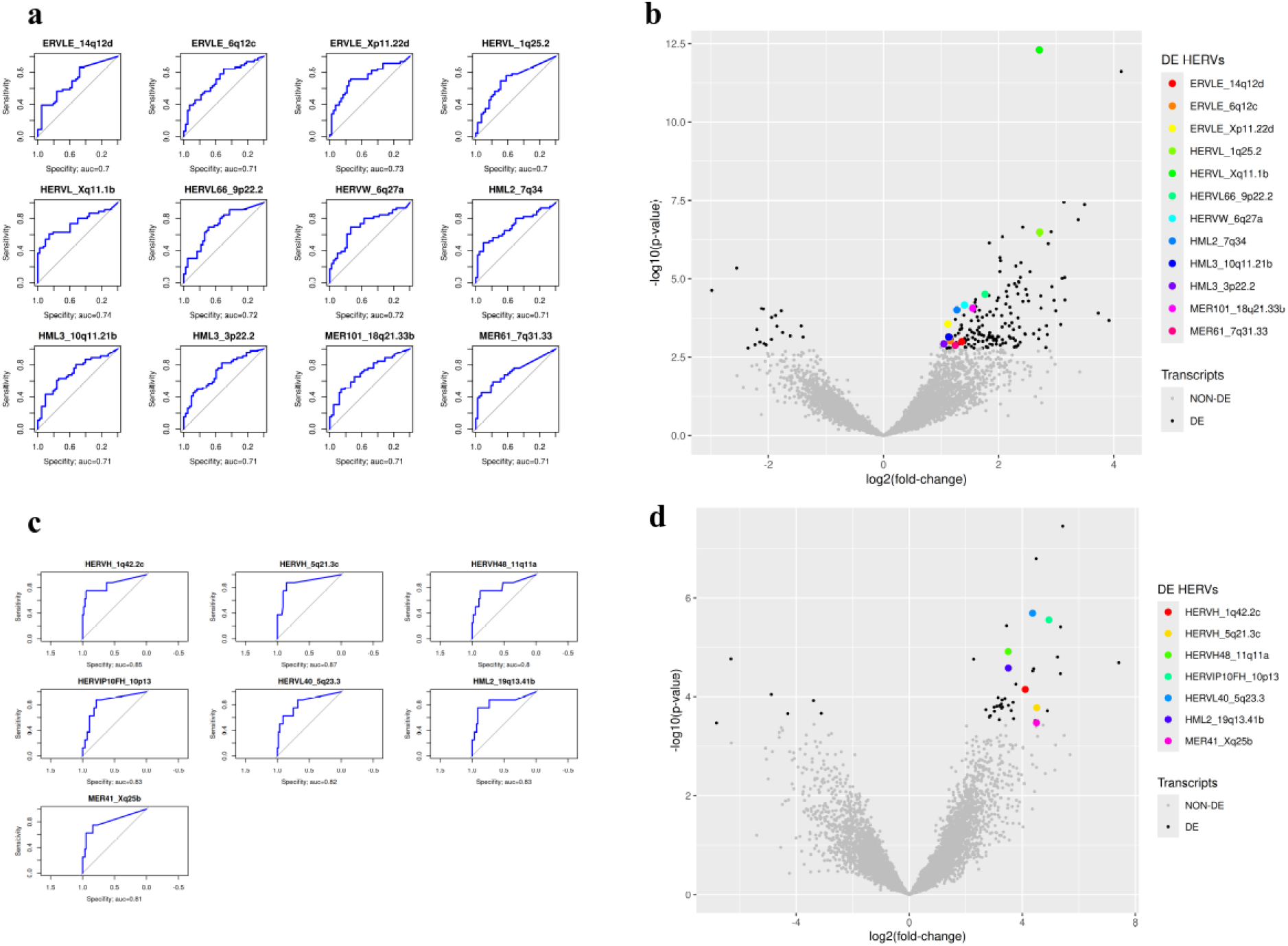
Differentially expressed HERVs for adenocarcinoma versus peritumoral and peritumoral (a, b) and for adenocarcinoma samples with *H. pylori* (c, d) using ROC curve and volcano plot, respectively. DE: differentially expressed; NON-DE: non differentially expressed; DE HERVs: Differentially expressed HERVs;

It is important to note, however, that the AUC values ranged from 0.70 to 0.74, indicating a degree of discriminative power considered reasonable. In samples with *H*.*pylori*, 7 HERVs belonging to the families HERVH, HERVH48, HERVIP10FH, HERVL40, HML2 and MER41 were found (Fig. 1c).

Among the HERVs identified by the ROC curve, ERV316A3_13q21.1a, ERVLE_6q12c, ERVLE_Xp11.22d, and HERVS71_3q12.3 showed higher expression, while ERVLB4_22q13.32b, HERV4_7q21.13a, HERVEA_19p13.3, HERVH_4q31.21c, HML3_3q22.1, and MER41_1q21.3b exhibited reduced expression trends, as illustrated in the volcano plot (Fig. 1b and 1d).

Subsequently, the volcano plot was used to assess which HERVs were being positively and negatively regulated. Next, the HERVs filtered based on the AUC metric were visualized in a heatmap. Tumor versus peritumoral tissue samples did not show clear clustering (Fig. 2a). In contrast, the heatmap comparing tumor tissues with and without *H. pylori* revealed higher HERV expression in *H. pylori*-positive samples, mainly HERVH48_11q11a and HERVH_1q42.2c (Fig. 2b).

**Fig. 2.**
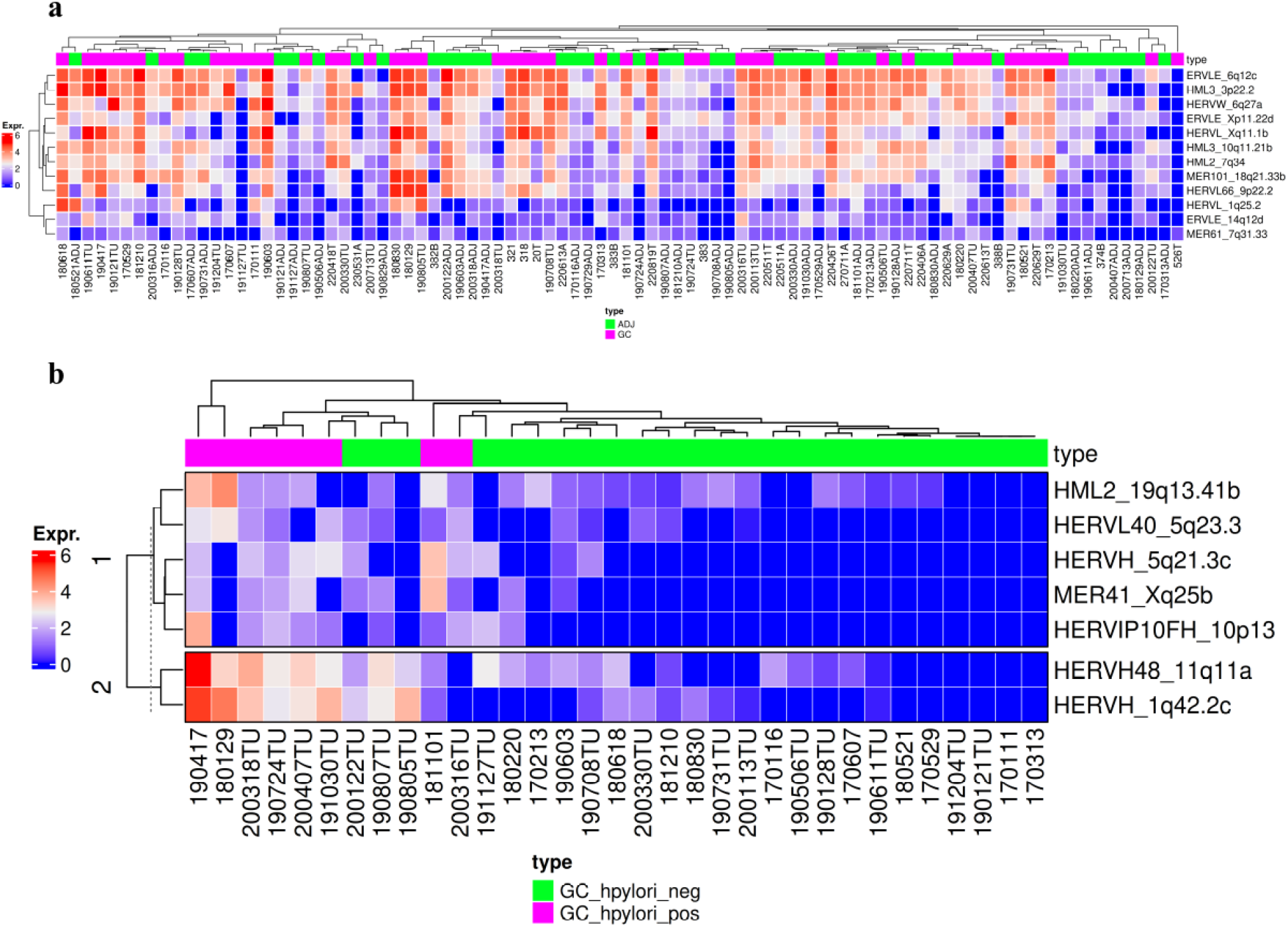
Heatmap of the expression patterns of differentially expressed HERVs for tumoral and peritumoral tissues (a) and tumoral tissues with *H. pylori* positive and negative (b). Expr.: Expression; ADJ: adjacent or peritumoral; GC: gastric cancer, or adenocarcinoma; GC_hpylori_ neg: *H. pylor*i-negative gastric cancer; GC_hpylori_ pos: *H. pylori*-positive gastric cancer.

After conducting statistical tests, it was observed that all HERVs present in the tumor tissue were upregulated compared to the peritumoral tissue (Fig. 3). Specifically, regarding tumor samples with *H. pylori* presence, those that tested positive exhibited increased HERV expression compared to the negative samples (Fig. 4).

**Fig. 3.**
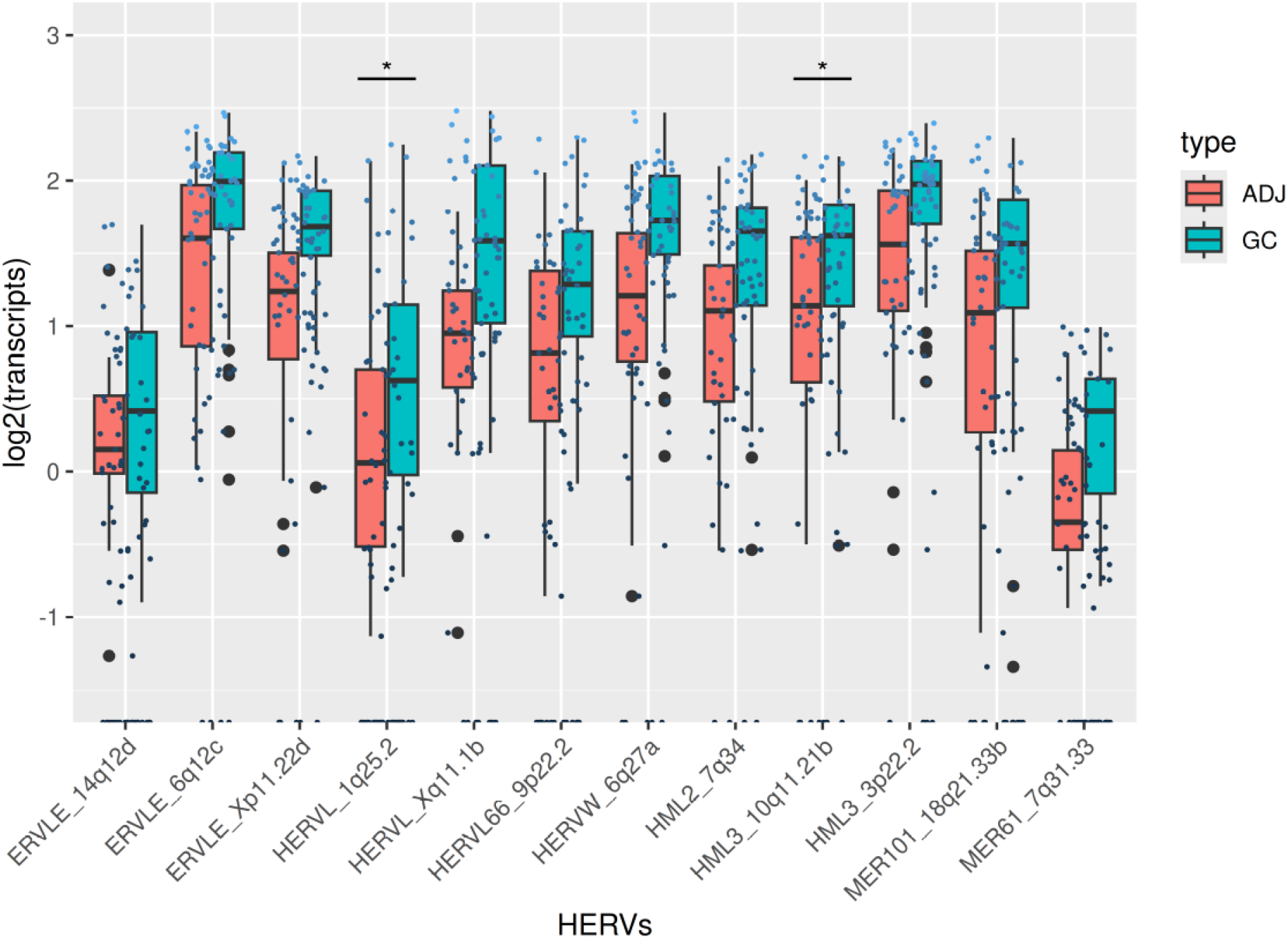
Boxplot of Differentially expressed HERVs for gastric cancer and adjacent tissues. p adj. < 0.0001. *: p. Adj. < 0.001.

**Fig. 4.**
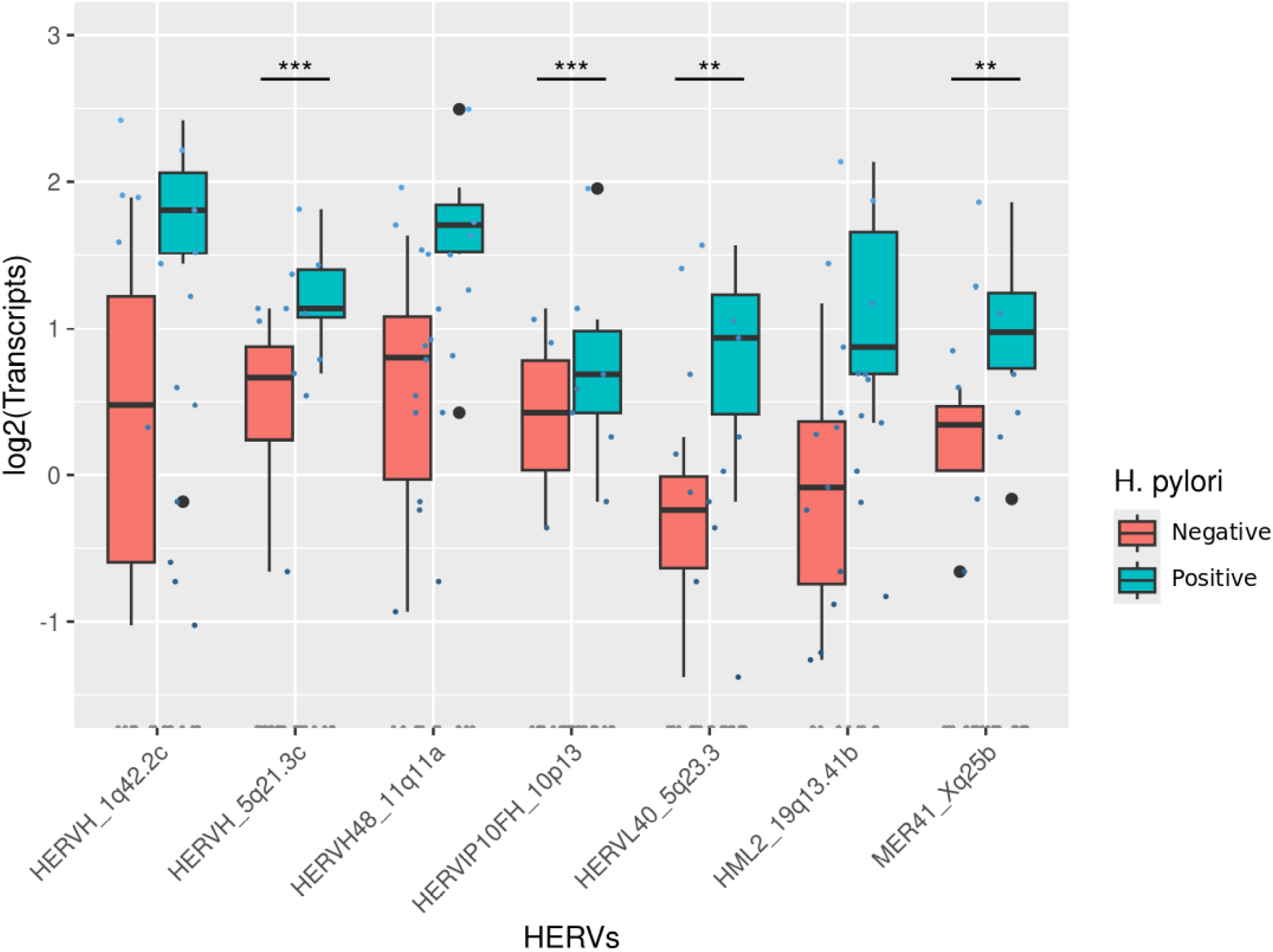
Boxplot of Differentially expressed HERVs for gastric cancer with *H. pylori* positive versus negative. p. adj. < 0.05. **: p. adj. < 0.01; ***: p. adj. < 0.001.

## DISCUSSION

In our study, differentially expressed HERVs were identified in gastric cancer. Although there is growing interest in HERVs within the academic community focused on malignancies, studies specifically addressing their role in gastric cancer remain scarce. Even rarer are investigations that explore the relationship between *H. pylori* and HERVs, and to date, no studies have integrated *H. pylori*, HERVs, and gastric cancer within a single framework.

A positive regulation of all relevant HERVs was observed in tumor samples, both in comparison to peritumoral tissues and in *H. pylori*-positive tumor samples. One possible explanation for this pattern is the demethylation of long terminal repeats (LTRs), which possess regulatory features that enable transcription by the cellular machinery, acting as long-range promoters or enhancers of host genes (Jern and Coffin, 2008). Conversely, host organisms have repressive mechanisms that recognize HERVs and regulate their activation (Geis and Goff, 2020).

HERVs can act as promoters or enhancers, providing transcription factor binding sites and regulatory signals that are recognized by RNA Polymerase II (Cherkasova, Chen and Childs, 2024; Jarosz and Halo, 2024). The silencing of HERVs occurs through histone modifications and DNA methylation. The primer binding site (PBS) serves as a repression mechanism during reverse transcription, mediated by members of the Krüppel-associated box zinc finger protein (KZFP) family, which form epigenetic silencing complexes and promote the spread of heterochromatin (Jarosz and Halo, 2024).

Normally, HERVs are silenced or exhibit marginal activity under basal conditions (Cortesi *et al*., 2024). The HUSH complex is one of the main agents responsible for this silencing, recruiting the chromatin remodeler MORC2 and the methyltransferase SETDB1 to promote the deposition of the epigenetic mark H3K9me3, thereby repressing HERV activity (Spencley *et al*., 2023). KZFP proteins also contribute to HERV silencing through mechanisms independent of the H3K9me3 mark (Xu *et al*., 2022).

However, this silencing can be reversed through hypomethylation and the loss of repressive histone marks, resulting in aberrant HERV expression (Krug et al., 2019). Furthermore, increased HERV expression leads to the production of double-stranded RNA (dsRNA) derived from these elements, which in turn triggers an interferon (IFN) response (Cañadas et al., 2018). Interferons are cytokines that regulate the immune system through various signaling pathways, contributing to cancer control by influencing the quality of antitumor immunity and enhancing responses to immunotherapy treatments (Spada and Ganguly, 2024).

The identification of double-stranded RNA (dsRNA) derived from HERVs by retroviral sensors in the cytoplasm is interpreted as a sign of viral infection, inducing a state of “viral mimicry” that triggers the production of type I and II interferons (Chiappinelli *et al*., 2015; Roulois *et al*., 2015). Viral mimicry is a process through which cells activate an innate immune response, inducing the expression of antiviral genes and potentially novel antigens (Jones *et al*., 2019).

This phenomenon holds therapeutic relevance in oncology, as HERVs may sensitize tumors to immunotherapy, enhancing the effectiveness of antitumor responses (Jansz and Faulkner, 2021).

Some HERVs reported in this study are associated with other types of cancer. In the case of tumoral and peritumoral analyses, the HML3_3p22.2 locus, for example, belongs to the HERVK9 family, also known as HERVK HML3, and contains the gag, pro, pol, and env genes (Mayer and Meese, 2002). This locus harbors a HERVK9 integration flanked by two solo LTRs known as MER9B (Kulski *et al*., 2008). Differential expression of HERVK9 has been reported in cervical cancer among racial subgroups of American women, Black and White, infected with HPV (Alldredge *et al*., 2023). Furthermore, HERVK9 may represent a common structural polymorphism within the MHC class I region (Stewart *et al*., 2004). In ovarian cancer patients, loss of HERVK9 associated with the epidermal growth factor receptor gene (EGFR) may be linked to prolonged survival in cases treated with platinum-based adjuvant chemotherapy (Fromhage *et al*., 2024)

The MER61_7q31.33 locus belongs to the HERV MER61 (Medium reiteration frequency interspersed repeat) family, a class I HERV flanked by the MER61 LTR, whose PBS shows similarity to that of HERVK9. The MER61 family is enriched with copies containing p53 binding sites, with a high likelihood of influencing gene regulation, either by direct enhancement or through cooperative action with other regulatory elements (Wang *et al*., 2007).

## CONCLUSION

Human endogenous retroviruses (HERVs) are viral DNA sequences that have integrated into the human genome over time and have lost their pathogenic potential. Recently, studies have investigated the role of these dysregulated elements in the development of cancer.

In this context, the present study identified 183 HERVs that were differentially expressed in tumor tissues compared to peritumoral tissues, as well as 44 and 104 HERVs associated with tumor tissues, depending on the presence of *H. pylori* infection. Among the transcripts described, 12 showed an AUC > 0.7 for the comparison between tumor and adjacent tissue, while 7 had an AUC > 0.8 for *H. pylori*. Our work is the first study to establish a relationship between differentially expressed HERVs, gastric cancer, and *H. pylori* infection. There is strong evidence that these HERVs may be regulated by hyper- or hypomethylated LTRs, potentially contributing to the development of gastric cancer. Thus, the HERVs identified in this study emerge as potential biomarkers for early diagnosis, prognosis, and monitoring of treatment response in patients with gastric cancer, including those with *H. pylori* infection.

## AUTHOR CONTRIBUTIONS

**Conceptualization**: Marcos da Conceição, Fabiano Cordeiro Moreira, Paulo Pimentel de Assumpção;

**Data curation**: Marcos da Conceição, Diego Pereira, Daniel de Souza Avelar, Fabiano Cordeiro Moreira;

**Formal analysis**: Marcos da Conceição, Sérgio Augusto Antunes Ramos, Diego Pereira, Ronald Matheus da Silva Mourão, Fabiano Cordeiro Moreira;

**Funding acquisition**: Fabiano Cordeiro Moreira, Geraldo Ishak, Samia Demachki, Samir Mansour Casseb, Paulo Pimentel de Assumpção, Rommel Mario Rodriguez Burbano, Williams Fernandes Barra;

**Investigation**: Marcos da Conceição, Fabiano Cordeiro Moreira; Samir Mansour Casseb;

**Methodology**: Marcos da Conceição, Juliana Barreto Albuquerque Pinto, Diego Pereira, Daniel de Souza Avelar, Jéssica Manoelli Costa da Silva, Ronald Matheus da Silva Mourão, Valéria Cristiane Santos da Silva, Rubem Ferreira da Silva, Amanda Ferreira Vidal; Tatiane Neotti, Ana Karyssa Mendes Anaissi, Samia Demachki, Samir Mansour Casseb, Fabiano Cordeiro Moreira;

**Sample Acquisition**: Geraldo Ishak, Paulo Pimentel de Assumpção, Rommel Mario Rodriguez Burbano, Williams Fernandes Barra;

**Supervision**: Fabiano Cordeiro Moreira, Samir Mansour Casseb, Paulo Pimentel de Assumpção;

**Project Administration**: Fabiano Cordeiro Moreira, Samir Mansour Casseb, Paulo Pimentel de Assumpção;

**Validation**: Tatiane Neotti, Ana Karyssa Mendes Anaissi, Samia Demachki, Williams Fernandes Barra;

**Visualization**: Marcos da Conceição, Sérgio Augusto Antunes Ramos,

**Writing – original draft**: Marcos da Conceição, Juliana Barreto Albuquerque Pinto, Valéria Cristiane Santos da Silva, Fabiano Cordeiro Moreira;

**Writing – review & editing**: Marcos da Conceição, Juliana Barreto Albuquerque Pinto, Valéria Cristiane Santos da Silva, Diego Pereira, Daniel de Souza Avelar, Jéssica Manoelli Costa da Silva, Ronald Matheus da Silva Mourão, Rubem Ferreira da Silva, Samir Mansour Casseb, Paulo Pimentel de Assumpção, Fabiano Cordeiro Moreira.

## REFERENCE

Alldredge, J. et al. (2023) “Endogenous Retrovirus RNA Expression Differences between Race, Stage and HPV Status Offer Improved Prognostication among Women with Cervical Cancer,” International Journal of Molecular Sciences, 24(2), p. 1492. Available at: 10.3390/ijms24021492.

Balakrishnan, M. et al. (2017) “Changing Trends in Stomach Cancer Throughout the World,” Current Gastroenterology Reports, 19(8), p. 36. Available at: 10.1007/s11894-017-0575-8.

Bao, C. et al. (2024) “Human endogenous retroviruses and exogenous viral infections,” Frontiers in Cellular and Infection Microbiology, 14, p. 1439292. Available at: 10.3389/fcimb.2024.1439292.

Bendall, M.L. et al. (2019) “Telescope: Characterization of the retrotranscriptome by accurate estimation of transposable element expression,” PLOS Computational Biology, 15(9), p. e1006453. Available at: 10.1371/journal.pcbi.1006453.

Bray, F. et al. (2024) Global cancer statistics 2022: GLOBOCAN estimates of incidence and mortality worldwide for 36 cancers in 185 countries - Bray - 2024 - CA: A Cancer Journal for Clinicians - Wiley Online Library. Available at: https://acsjournals.onlinelibrary.wiley.com/doi/10.3322/caac.21834 (Accessed: September 4, 2025).

Burn, A. et al. (2022) “Widespread expression of the ancient HERV-K (HML-2) provirus group in normal human tissues,” PLoS Biology, 20(10), p. e3001826. Available at: 10.1371/journal.pbio.3001826.

Cañadas, I. et al. (2018) “Tumor innate immunity primed by specific interferon-stimulated endogenous retroviruses,” Nature medicine, 24(8), pp. 1143–1150. Available at: 10.1038/s41591-018-0116-5.

Cegolon, L. et al. (2013) “Human endogenous retroviruses and cancer prevention: evidence and prospects,” BMC cancer, 13, p. 4. Available at: 10.1186/1471-2407-13-4.

Cherkasova, E.A., Chen, L. and Childs, R.W. (2024) “Mechanistic regulation of HERV activation in tumors and implications for translational research in oncology,” Frontiers in Cellular and Infection Microbiology, 14, p. 1358470. Available at: 10.3389/fcimb.2024.1358470.

Chiappinelli, K.B. et al. (2015) “Inhibiting DNA methylation causes an interferon response in cancer via dsRNA including endogenous retroviruses,” Cell, 162(5), pp. 974–986. Available at: 10.1016/j.cell.2015.07.011.

Cortesi, A. et al. (2024) “Activation of endogenous retroviruses and induction of viral mimicry by MEK1/2 inhibition in pancreatic cancer,” Science Advances, 10(13), p. eadk5386. Available at: 10.1126/sciadv.adk5386.

Curado, M.P. et al. (2019) “Disparities in Epidemiological Profile of Gastric Adenocarcinoma in Selected Cities of Brazil,” Asian Pacific Journal of Cancer Prevention: APJCP, 20(8), pp. 2253–2258. Available at: 10.31557/APJCP.2019.20.8.2253.

Dobin, A. et al. (2013) “STAR: ultrafast universal RNA-seq aligner,” Bioinformatics, 29(1), pp. 15–21. Available at: 10.1093/bioinformatics/bts635.

Duan, Y. et al. (2025) “Helicobacter pylori and gastric cancer: mechanisms and new perspectives,” Journal of Hematology & Oncology, 18, p. 10. Available at: 10.1186/s13045-024-01654-2.

Fromhage, G. et al. (2024) “Loss of copy numbers of retrotransposons (HERVK) on chromosome 7p11.2 impacts EGFR (Epidermal Growth Factor Receptor)-induced phenotypes for platinum sensitivity and long-term survival in ovarian cancer—A study from the OVCAD consortium,” International Journal of Cancer, 155(5), pp. 934–945. Available at: 10.1002/ijc.34976.

Geis, F.K. and Goff, S.P. (2020) “Silencing and Transcriptional Regulation of Endogenous Retroviruses: An Overview,” Viruses, 12(8), p. 884. Available at: 10.3390/v12080884.

Gu, Z. (2022) “Complex heatmap visualization,” iMeta, 1(3), p. e43. Available at: 10.1002/imt2.43.

Hooi, J.K.Y. et al. (2017) “Global Prevalence of Helicobacter pylori Infection: Systematic Review and Meta-Analysis,” Gastroenterology, 153(2), pp. 420–429. Available at: 10.1053/j.gastro.2017.04.022.

Hu, R. et al. (2024) “Short Read RNA-Seq,” Methods in molecular biology (Clifton, N.J.), 2822, pp. 245–262. Available at: 10.1007/978-1-0716-3918-4_17.

IARC (1994) “Infection with Helicobacter pylori,” IARC monographs on the evaluation of carcinogenic risks to humans, 61, pp. 177–240.

Jansz, N. and Faulkner, G.J. (2021) “Endogenous retroviruses in the origins and treatment of cancer,” Genome Biology, 22, p. 147. Available at: 10.1186/s13059-021-02357-4.

Jarosz, A.S. and Halo, J.V. (2024) “Transcription of Endogenous Retroviruses: Broad and Precise Mechanisms of Control,” Viruses, 16(8), p. 1312. Available at: 10.3390/v16081312.

Jern, P. and Coffin, J.M. (2008) “Effects of Retroviruses on Host Genome Function,” Annual Review of Genetics, 42(Volume 42, 2008), pp. 709–732. Available at: 10.1146/annurev.genet.42.110807.091501.

Johnson, W.E. (2019) “Origins and evolutionary consequences of ancient endogenous retroviruses,” Nature Reviews Microbiology, 17(6), pp. 355–370. Available at: 10.1038/s41579-019-0189-2.

Jones, P.A. et al. (2019) “Epigenetic therapy in immune-oncology,” Nature Reviews Cancer, 19(3), pp. 151–161. Available at: 10.1038/s41568-019-0109-9.

Kulski, J.K. et al. (2008) “Human Endogenous Retrovirus (HERVK9) Structural Polymorphism With Haplotypic HLA-A Allelic Associations,” Genetics, 180(1), pp. 445–457. Available at: 10.1534/genetics.108.090340.

Lerat, E. (2010) “Identifying repeats and transposable elements in sequenced genomes: how to find your way through the dense forest of programs,” Heredity, 104(6), pp. 520–533. Available at: 10.1038/hdy.2009.165.

Li, Y., He, C. and Lu, N. (2024) “Impacts of Helicobacter pylori infection and eradication on gastrointestinal microbiota: An up-to-date critical review and future perspectives,” Chinese Medical Journal, 137(23), pp. 2833–2842. Available at: 10.1097/CM9.0000000000003348.

Love, M.I., Huber, W. and Anders, S. (2014) “Moderated estimation of fold change and dispersion for RNA-seq data with DESeq2,” Genome Biology, 15(12), p. 550. Available at: 10.1186/s13059-014-0550-8.

Löwer, R. (1999) “The pathogenic potential of endogenous retroviruses: facts and fantasies,” Trends in Microbiology, 7(9), pp. 350–356. Available at: 10.1016/S0966-842X(99)01565-6.

Mayer, J. and Meese, E.U. (2002) “The Human Endogenous Retrovirus Family HERV-K(HML-3),” Genomics, 80(3), pp. 331–343. Available at: 10.1006/geno.2002.6839.

Plummer, M. et al. (2015) “Global burden of gastric cancer attributable to Helicobacter pylori,” International Journal of Cancer, 136(2), pp. 487–490. Available at: 10.1002/ijc.28999.

R Core Team (2025) “R: A Language and Environment for Statistical Computing.” Viena, Autria: R Foundation for Statistical Computing. Available at: https://www.R-project.org/ (Accessed: September 4, 2025).

Robin, X. et al. (2011) “pROC: an open-source package for R and S+ to analyze and compare ROC curves,” BMC Bioinformatics, 12(1), p. 77. Available at: 10.1186/1471-2105-12-77.

Roulois, D. et al. (2015) “DNA-demethylating agents target colorectal cancer cells by inducing viral mimicry by endogenous transcripts,” Cell, 162(5), pp. 961–973. Available at: 10.1016/j.cell.2015.07.056.

Spada, S. and Ganguly, A. (2024) “Chapter Four - Role of interferon dependent and independent signaling pathways: Implications in cancer,” in S. Mukherjee and K. Chatterjee (eds.) International Review of Cell and Molecular Biology. Academic Press (Targeting Signaling Pathways in Solid Tumors - Part C), pp. 153–162. Available at: 10.1016/bs.ircmb.2024.06.004.

Spencley, A.L. et al. (2023) “Co-transcriptional genome surveillance by HUSH is coupled to termination machinery,” Molecular Cell, 83(10), pp. 1623-1639.e8. Available at: 10.1016/j.molcel.2023.04.014.

Srinivasachar Badarinarayan, S. and Sauter, D. (2022) “Not all viruses cause disease: HERV-K(HML-2) in healthy human tissues,” PLOS Biology, 20(10), p. e3001884. Available at: 10.1371/journal.pbio.3001884.

Stewart, C.A. et al. (2004) “Complete MHC Haplotype Sequencing for Common Disease Gene Mapping,” Genome Research, 14(6), pp. 1176–1187. Available at: 10.1101/gr.2188104.

Vargiu, L. et al. (2016) “Classification and characterization of human endogenous retroviruses; mosaic forms are common,” Retrovirology, 13, p. 7. Available at: 10.1186/s12977-015-0232-y.

Wang, T. et al. (2007) “Species-specific endogenous retroviruses shape the transcriptional network of the human tumor suppressor protein p53,” Proceedings of the National Academy of Sciences of the United States of America, 104(47), pp. 18613–18618. Available at: 10.1073/pnas.0703637104.

Xu, R. et al. (2022) “Stage-specific H3K9me3 occupancy ensures retrotransposon silencing in human pre-implantation embryos,” Cell Stem Cell, 29(7), pp. 1051-1066.e8. Available at: 10.1016/j.stem.2022.06.001.

